# Preferred synonymous codons are translated more accurately: Proteomic evidence, among-species variation, and mechanistic basis

**DOI:** 10.1101/2022.02.22.481448

**Authors:** Mengyi Sun, Jianzhi Zhang

## Abstract

A commonly stated cause of the widespread phenomenon of unequal uses of synonymous codons is their differential translational accuracies. However, this long-standing translational accuracy hypothesis (TAH) of codon usage bias has had no direct evidence beyond anecdotes. Analyzing proteomic data from *Escherichia coli*, we observe higher translational accuracies of more frequently used synonymous codons, offering direct, global evidence for the TAH. The experimentally measured codon-specific translational accuracies validate a sequence-based proxy; this proxy provides support for the TAH from the vast majority of over 1000 taxa surveyed in all domains of life. We find that the relative translational accuracies of synonymous codons vary substantially among taxa and are strongly correlated with the amounts of cognate tRNAs relative to those of near-cognate tRNAs. These and other observations suggest a model in which selections for translational efficiency and accuracy drive codon usage bias and its coevolution with the tRNA pool.

## INTRODUCTION

Eighteen of the 20 amino acids are each encoded by more than one codon, but the synonymous codons are usually unequally used in a genome (*1, 2*). Among the synonymous codons of an amino acid, those used more often than the average are referred to as preferred codons while the rest unpreferred. This phenomenon of codon usage bias (CUB), initially discovered over four decades ago from the first few determined gene sequences (*3-6*), is a result of the joint forces of mutation, genetic drift, and natural selection, but the specific selective agents have not been fully deciphered (*1, 2*). One long-standing hypothesis known as the translational accuracy hypothesis (TAH) asserts that different synonymous codons are translated with different accuracies and that CUB results at least in part from natural selection for translational accuracy (*7*). Indeed, the importance of accurate protein translation cannot be overstated, because mistranslation may lead to the loss of normal protein functions and gain of cellular toxicity (*8*) and cause severe diseases including cancer and neurodegenerative diseases (*9*). In fact, several cellular mechanisms are known to ensure the overall fidelity of protein synthesis. For example, conformational changes of the ribosome decoding center can be more efficiently induced by cognate codon-anticodon interactions than near-cognate codon-anticodon interaction (*10*), allowing discrimination against incorrect decoding. Additionally, the accuracy of many steps in translation, such as tRNA aminoacylation (*10*) and codon-anticodon matching, is enhanced by the energy-consuming kinetic proofreading (*11*). Notwithstanding, even if synonymous codons differ in translational accuracy, relatively accurate synonymous codons may not be preferentially used. This is because synonymous codons differ in other properties such as the translational elongation speed (*12, 13*); selection related to these other features(*14*) could triumph selection for translational accuracy.

Several groups have attempted to test the TAH of CUB. In particular, Akashi (*7*) developed an indirect test based on the idea that the benefit of using relatively accurate codons should be greater at evolutionarily conserved amino acid sites than unconserved sites of the same protein; consequently, a higher usage of preferred codons at conserved sites than at unconserved sites supports the TAH. While Akashi’s test is positive for several model organisms investigated (*7, 15, 16*), this test does not directly compare translational accuracies of synonymous codons so cannot completely exclude alternative explanations (*7, 17*). In an early study, Precup and Parker used site-directed mutagenesis followed by peptide sequencing to show that AAU, an unpreferred codon of Asn, is misread as Lys 4-9 times more often than is AAC, a preferred codon of Asn, at a particular position of the coat protein gene of the bacteriophage MS2 under Asn starvation (*18*). Similarly, Kramer and Farabaugh observed that AAU has a significantly higher rate of mistranslation to Lys than AAC at a particular position of a reporter gene in *Escherichia coli* (*19*). Nonetheless, Kramer and Farabaugh also observed that the unpreferred Arg codons of CGA and CGG and the preferred Arg codons of CGU and CGC exhibited similar rates of mistranslation to Lys (*19*). While the above experiments directly tested the TAH, they were each based on the investigation of one amino acid site of one protein, so its genome-wide generality is unknown. As such, a direct and global test of the TAH is needed.

Capitalizing on a proteome-wide probe of mistranslation in *E. coli* (*20*), we here provide direct evidence that preferred codons are generally translated more accurately than unpreferred codons. We then use the *E. coli* data to validate a sequence-based proxy for relative translational accuracies of synonymous codons. Using this proxy, we show that the TAH of CUB is supported in the vast majority of over 1000 diverse taxa surveyed, but that the relative translational accuracies of synonymous codons vary substantially among taxa. We find that the relative translational accuracy of a synonymous codon is strongly correlated with its cognate tRNA abundance relative to near-cognate tRNA abundance. These and other results suggest a model in which selections for translational efficiency and accuracy drive the CUB and its coevolution with the tRNA pool.

## RESULTS

### Preferred codons are more accurately decoded

A direct test of the TAH of CUB requires comparing the mistranslation rate among synonymous codons. Using mass spectrometry, Mordret *et al*. quantified mistranslations at individual sites of the *E. coli* proteome (*20*). After removing sites and codons where mistranslation rates cannot be quantified due to technical reasons (see Materials and Methods), we grouped mistranslation events according to the identities of their original codons. We then computed the absolute mistranslation rate of a codon as the ratio of the total intensity of mistranslated peptides to that of all peptides mapped to the codon. Finally, we computed the relative mistranslation rate (*RMR*) of a codon by dividing its absolute mistranslation rate by the mean absolute mistranslation rate of all codons coding for the same amino acid. *RMR* >1 means that the codon has a higher mistranslation rate than the average among all codons for the same amino acid, whereas *RMR* <1 means the opposite. Codon usage was assessed by the relative synonymous codon usage (*RSCU*). The *RSCU* of a codon equals its frequency in the genome relative to the average frequency of all codons for the same amino acid (*21*). A codon with *RSCU* >1 is preferred while a codon with *RSCU* <1 is unprefered.

We were able to estimate the *RMR* for 27 codons of nine amino acids (**Fig. 1a**). Except for Gly, the most preferred synonymous codon of an amino acid shows *RMR* <1, providing a significant support for the TAH (*P* = 0.020, one-tailed binomial test). Similarly, except for Gly and Val, the least prevalent synonymous codon of an amino acid shows *RMR* >1 (*P* = 0.090, one-tailed binomial test). Because both *RSCU* and *RMR* of a codon are relative to the mean of all codons for the same amino acid, they can be compared among codons of different amino acids. Indeed, a strong negative correlation was observed between *RSCU* and *RMR* among the 27 codons (Pearson’s *r* = -0.56, *P* < 0.001, permutation test; Spearman’s *ρ* = -0.49, *P* = 0.005, permutation test; **Fig. 1b**). Together, these findings from the proteomic data of *E. coli* demonstrate that preferred codons tend to have lower mistranslation rates, supporting the TAH of CUB.

**Figure 1.**
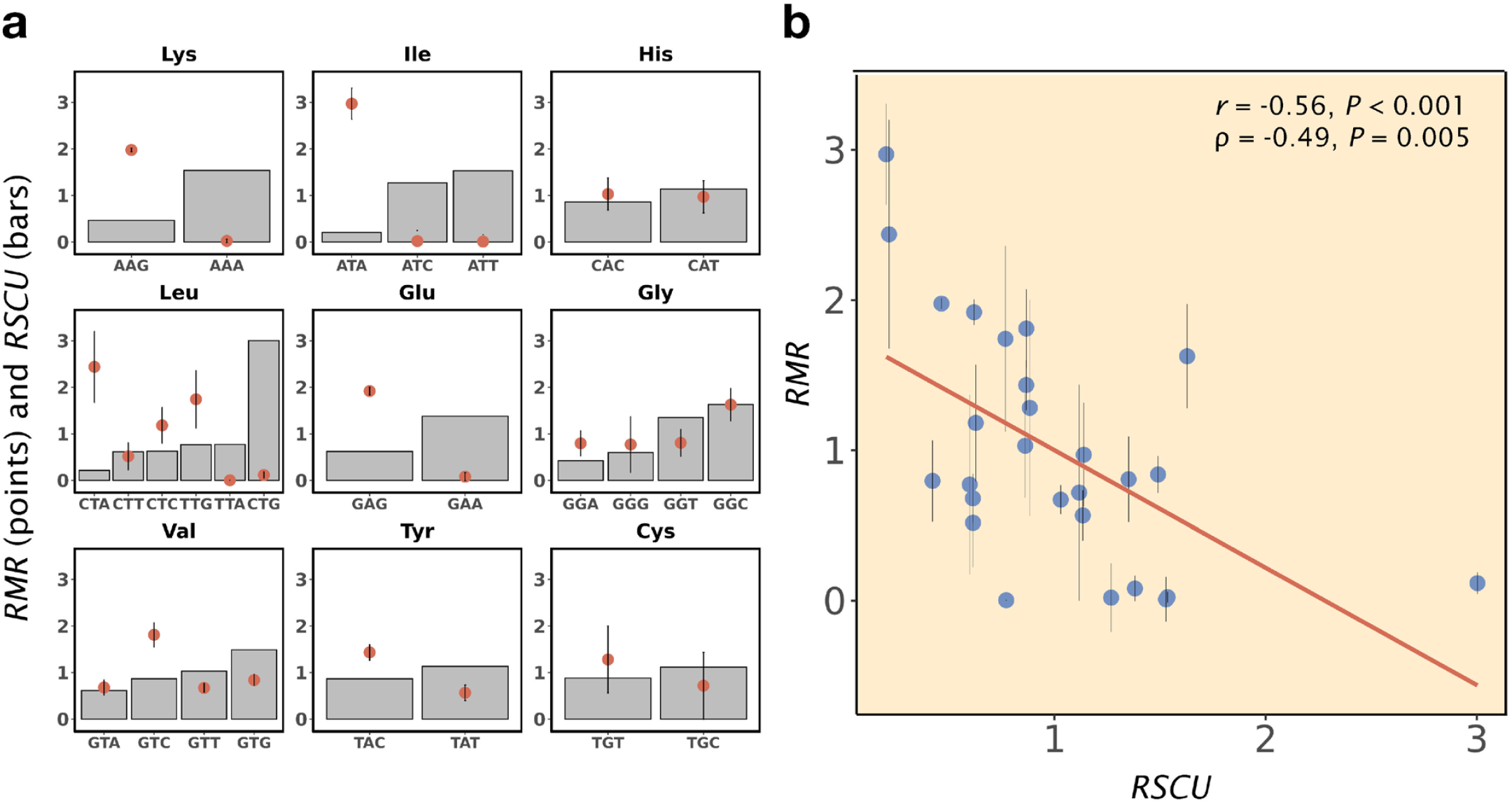
More frequently used synonymous codons tend to be decoded more accurately in *E. coli*. **a**, Comparison of relative synonymous codon usage (*RSCU*, bars) and relative mistranslation rate (*RMR*, dots) among synonymous codons for nine amino acids with proteome-based *RMR* estimates. **b**, A significant negative correlation between *RSCU* and *RMR* across the 27 codons in panel a. The red line is the linear regression. In both panels, error bars represent one standard error estimated by the bootstrap method. The standard error of *RSCU* estimated by the bootstrap method is negligible due to the large number of each codon in the genome, so is not shown. *P*-values are based on permutation tests.

### Relative translational accuracies of synonymous codons vary across taxa

How do certain synonymous codons achieve higher translational accuracies than others? There are two general scenarios. In the first scenario, referred hereinafter as the constant accuracy hypothesis, the translational accuracy is intrinsically higher for a synonymous codon than another because of their different chemical nature (*22*). Consequently, the relative translational accuracies of synonymous codons should be more or less the same in different species. For instance, given that AAA (Lys) is more accurate than AAG (Lys) in *E. coli* (**Fig. 1a**), we expect the same trend in the vast majority if not all species. Alternatively, relative translational accuracies of synonymous codons may be greatly influenced by species-specific factors such as the tRNA pool. Under this scenario, referred to as the variable accuracy hypothesis hereinafter, the relative accuracies of synonymous codons vary among species. That is, AAA is more accurate than AAG in many species but the opposite is true in many other species.

Measuring the relative translational accuracies of synonymous codons in a large number of species will allow differentiating between the above two hypotheses, which will in turn help understand the mechanism underlying the translational accuracy differences among synonymous codons. Because codon-specific, proteome-based translational accuracies have not been measured beyond *E. coli*, we resort to a sequence-based proxy referred to as the odds ratio (*OR*) that originated from Akashi’s test (*7*). Specifically, the *OR* of synonymous codon X that encodes amino acid Y in a gene is the number of times that X is used at invariant Y sites relative to the number of times that X is not used at invariant Y sites, divided by the number of times that X is used at variant Y sites relative to the number of times that X is not used at variant Y sites (**Fig. 2a**). Here, invariant and variant Y sites refer to Y sites in the focal species whose counterparts in the ortholog from a related species have Y and non-Y, respectively. The *OR* values computed from individual genes can be combined to yield a single *OR* using the Mantel-Haenszel procedure (see Materials and Methods). While *OR* was originally developed for preferred codons, it can be computed for any codon of the 18 amino acids that have multiple synonymous codons (*14*). Based on Akashi’s test, *OR* has been used as a proxy for the relative translational accuracy of a codon (*14*). To verify the relationship between *OR* and relative translational accuracy, we computed *OR* values by aligning *E. coli* genes with their *Salmonella enterica* orthologs. Indeed, for the 27 codons with *RMR* estimates, *OR* and *RMR* are strongly negatively correlated (*r* = -0.63, *P* = 0.001; *ρ* = -0.43, *P* = 0.01; **Fig. 2b**), confirming that the *OR* of a codon is a valid proxy for its relative translational accuracy.

**Figure 2.**
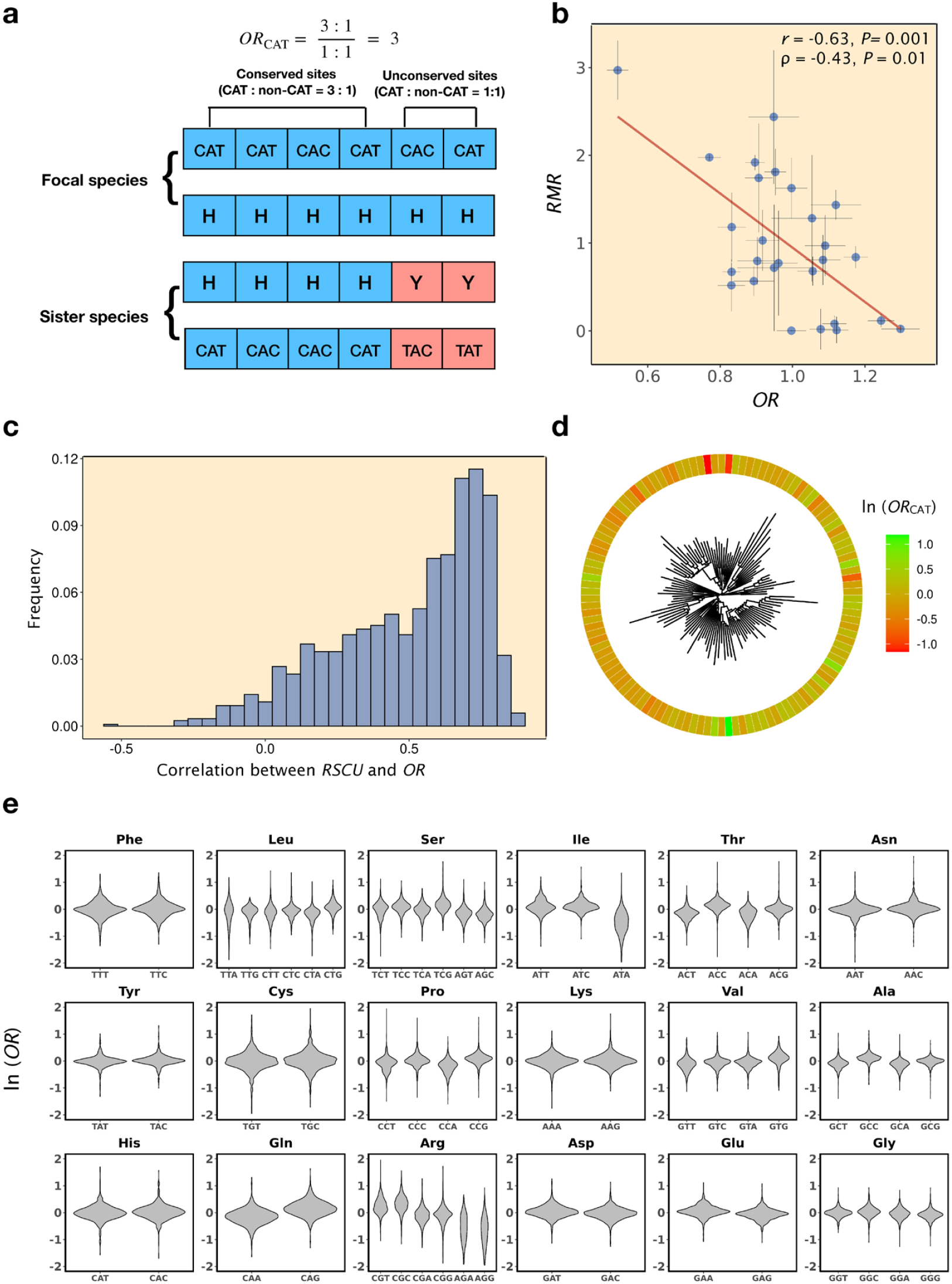
Variation of relative translational accuracies of synonymous codons across taxa. **a**, Diagram explaining the calculation of odds ratio (*OR*) of the codon CAT that serves as a proxy for its relative translational accuracy. Showing here is a hypothetical alignment of orthologous proteins (and the underlying coding sequences) between the focal species and a related species. **b**, *OR* is negatively correlated with *RMR* across codons in *E. coli. P*-values are based on permutation tests. The red line is the linear regression. **c**, Frequency distribution of Pearson’s correlation between *RSCU* and *OR* in 1197 bacterial taxa. Ninety-five percent of these taxa show positive correlations. **d**, ln(*OR*) of codon CAT for each of 118 bacterial taxa, one per order, arranged according to their phylogeny shown in the middle. **e**, Violin plots showing frequency distributions of ln(*OR*) of individual codons of the 18 amino acids that have multiple synonymous codons across 1197 bacterial taxa.

To examine whether the relative translational accuracies of synonymous codons vary among species, we took advantage of a recently built phylogenetic tree of 10,575 microbial taxa (*23*). Because most taxa (9,906) in the tree are from the domain Bacteria, we first focused our analysis on Bacteria. We picked all 1,197 pairs of sister bacterial taxa from the tree and aligned their orthologous genes (see Materials and Methods). We randomly assigned one taxon in each pair as the focal taxon and computed *OR* for each codon as described above. We found a positive correlation between *RSCU* and *OR* across codons in 95% of the taxa examined (**Fig. 2c**), demonstrating an overwhelming support for the TAH of CUB in Bacteria.

We computed ln(*OR*) to make its distribution relatively symmetric to aid visualization, and examined as an example ln(*OR*) for codon CAT (His) in each of the focal taxa arranged according to the bacterial tree (one taxon per order is presented in **Fig. 2d**). We found ln(*OR*) to vary greatly from negative values to positive values, with a high density near 0 (**Fig. 2e**). Furthermore, the extreme values of ln(*OR*) (bright red and bright green in **Fig. 2d**) are scattered across the tree rather than concentrated in a few clades, suggesting that the relative translational accuracy of CAT has changed substantially and frequently in evolution. The across-taxon variation of *OR* indicates that CAT is the relatively inaccurate one of the two synonymous codons of His in many taxa (red in **Fig. 2d**) but the relatively accurate one in many other taxa (green), supporting the variable accuracy hypothesis. From **Fig. 2e**, which shows the 18 amino acids each with multiple codons, it is clear that the pattern observed for CAT applies to all codons. Furthermore, every codon has *OR* >1 in at least 8.9% of the taxa examined (**Fig. S1a**). These results thus support the variable accuracy hypothesis for all synonymous codons. The above observations of *OR* variation among taxa are not primarily caused by sampling error, because a similar pattern was detected when we analyzed a subset of taxa for each amino acid where the number of occurrences of each synonymous codon considered in *OR* estimation is at least 1000 per taxon (**Fig. S1b**). They are not mainly caused by genetic drift either, because a similar pattern was found when we analyzed a subset of taxa with strong signals of selection for translational accuracy (correlation between *RSCU* and *OR* exceeding 0.5) (**Fig. S1c**). It is worth pointing out that, despite the general support for the variable accuracy hypothesis, for a minority of codons such as ATA (Ile), AGA (Arg), and AGG (Arg), the distribution of ln(*OR*) is strongly skewed toward negative values (**Fig. 2e**), suggesting that their relative translational accuracies are somewhat constrained although not invariable in evolution.

To investigate if the above observations from Bacteria are generalizable to the other two domains of life, we first expanded our analysis to Archaea represented in the large phylogeny mentioned (*23*). We found that the correlation between *RSCU* and *OR* is positive in 90% of taxa examined and that ln(*OR*) varies greatly across taxa for each codon (**Fig. S2**), further supporting the TAH and the variable accuracy hypothesis. For Eukaryota, we analyzed five commonly used model organisms: human, mouse, worm, fly, and budding yeast (see Materials and Methods). In each of these species, the correlation between *RSCU* and *OR* is significantly positive (**Table S1**), supporting the TAH. Except for the two mammals, which are closely related, the *OR*s estimated from one species are not well correlated with those estimated from another species (**Fig. S3**). Furthermore, the correlation in *OR* generally declines with the divergence time between the two species (**Fig. S3**), consistent with the variable accuracy hypothesis. Taken together, our results show that the TAH is generally supported in all domains of life but the relative translational accuracies of synonymous codons vary across taxa.

### Mechanistic basis of among-codon and across-taxon variations of translational accuracies

The empirical support for the variable accuracy hypothesis strongly suggests that the determinants of the *RMR*s of synonymous codons vary among species. In the aforementioned Kramer-Farabaugh study (*19*), the authors found that artificially increasing the expression level of the cognate tRNA for Arg codons AGA and AGG reduces their mistranslations to Lys, so proposed that the competition between cognate and near-cognate tRNAs determines the mistranslation rate of a codon. Here, the cognate tRNA is the tRNA whose anticodon pairs with the codon correctly (allowing wobble pairing), whereas the near-cognate tRNA corresponds to a different amino acid and has an anticodon that mismatches the codon at one position. Consistent with the above proposal, Mordret *et al*. (*20*) inferred that most of the mistranslation events in *E. coli* arose from mispairing between codons and near-cognate tRNAs. They further noted that, for certain types of mistranslation, there is a negative correlation across growth phases between the mistranslation rate and the ratio (*R*_c/nc_) in abundance between cognate and near-cognate tRNAs, although the correlation was rarely statistically significant (*20*). Based on these past observations, we hypothesize that the relative translational accuracy of a synonymous codon increases with its relative *R*_c/nc_, or *RR*_c/nc_, which is *R*_c/nc_ divided by the mean *R*_c/nc_ of all codons coding for the same amino acid (see Materials and Methods). We further hypothesize that, because the tRNA pool varies substantially among species (*24*), the among-species variation of relative translational accuracies arises from the among-species variation in *RR*_c/nc_.

To test the above hypotheses, we computed *RR*_c/nc_ for each codon using published tRNA expression levels in *E. coli* (*20*). Indeed, we observed a significant negative correlation between *RR*_c/nc_ and *RMR* (*r* = -0.47, *P* = 0.009; *ρ* = -0.53, *P* = 0.005; **Fig. 3a**) and a significant positive correlation between *RR*_c/nc_ and *OR* (*r* = 0.49, *P* = 0.07; *ρ* = 0.74, *P* = 0.00001; **Fig. 3b**) across codons, supporting the hypothesis that the relative ratio of cognate to near-cognate tRNA abundances is a major determinant of a codon’s relative translational accuracy in *E. coli*. Note that the relative cognate tRNA abundance alone is not significantly correlated with *RMR* (*r* = - 0.22, *P* = 0.2; ρ = -0.04, *P* = 0.4; **Fig. S4a**), supporting the role of competition between cognate and near-cognate tRNAs in determining *RMR*. As previously reported (*14*), the relative cognate tRNA level is highly correlated with *RSCU* (*r* = 0.61, *P* = 0.03; *ρ* = 0.48, *P* = 0.02; **Fig. S4b**), which is likely a result of selection for high translational efficiency (i.e., more codons translated per unit time per cell) because balanced codon usage relative to cognate tRNA concentrations maximizes translational efficiency (*14*).

**Figure 3.**
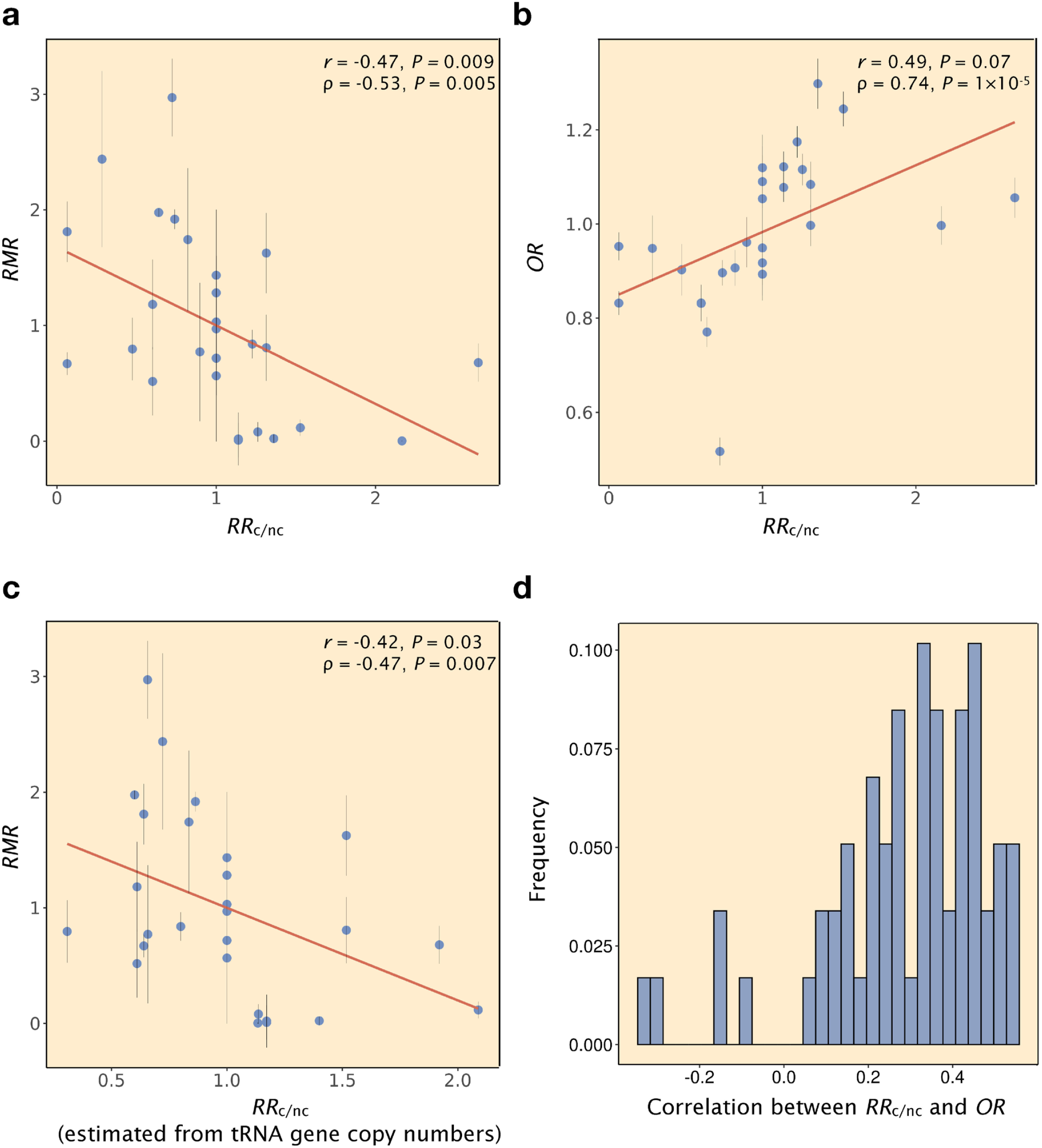
The relative ratio of cognate tRNA concentration to near-cognate tRNA concentration (*RR*_c/nc_) is a major determinant of a codon’s relative translational accuracy. **a**, *RMR* is negatively correlated with *RR*_c/nc_ across codons in *E. coli*. **b**, *OR* is positively correlated with *RR*_c/nc_ across codons in *E. coli*. **c**, *RMR* is negatively correlated with *RR*_c/nc_ computed using tRNA gene copy numbers instead of tRNA concentrations in *E. coli*. **d**, Frequency distribution of Pearson’s correlation between *OR* and *RR*_c/nc_ computed using tRNA gene copy numbers in bacterial taxa with >80 tRNA genes. All *P*-values are based on permutation tests. In a-c, the red line is the linear regression.

We next investigated whether the above finding in *E. coli* applies to other bacterial taxa surveyed in Fig. 2. Because tRNA expression levels are unknown for the vast majority of these taxa, we used the gene copy number of each tRNA species as a proxy for the total expression level of the tRNA species (*25*). Indeed, *E. coli RR*_c/nc_ computed from tRNA gene copy numbers is highly correlated with that computed from tRNA expression levels (*r* = 0.77, *P* = 1.64×10^−6^; *ρ* = 0.90, *P* = 1.36×10^−10^). Furthermore, *E. coli RR*_c/nc_ computed from tRNA gene copy numbers is significantly correlated with *RMR* (**Fig. 3c**), confirming the validity of using this proxy. We obtained the tRNA gene annotations for 1094 of the 1197 focal bacterial taxa examined in Fig. 2. However, in many of these taxa, there is little tRNA gene redundancy or variation in cognate tRNA gene copy number among synonymous codons despite considerable CUB; in these taxa, the tRNA gene copy number is unlikely a good proxy for tRNA abunadnce (*26*). Because the tRNA gene copy number is a good proxy for tRNA abundance in *E. coli*, which has 85 tRNA genes, we decided to filter out taxa with fewer than 81 tRNA genes to strike a balance between the noise level and number of taxa in our analysis. This filtering left us with 59 taxa, in each of which we correlated the *OR* of a codon with its *RR*_c/nc_ computed from tRNA gene copy numbers. The vast majority (92%) of the taxa show a positive correlation (**Fig. 3d**), supporting the generality of our hypothesis on the role of *RR*_c/nc_ in determining the relative translational accuracy of a codon in Bacteria.

To investigate whether the above finding is generalizable to other domains of life, we analyzed tRNA genes in Archaea taxa and Eukaryotic model organisms. Unfortunately, no Archaea taxa examined have more than 80 tRNA genes. For each of the five eukaryotes (human, mouse, fly, worm, and yeast), the correlation between *OR* and *RR*_c/nc_ computed from tRNA gene copy numbers is significantly positive for linear or rank correlation (**Table S2**). Together, our findings strongly support that, in the diverse taxa surveyed, the ratio of cognate tRNA abundance to near-cognate tRNA abundance is generally a major determinant of the relative translational accuracy of a codon. Hence, the variation of the tRNA pool among species can explain the across-species variation of the relative translational accuracies of synonymous codons.

## DISCUSSION

Analyzing published proteomic data from *E. coli*, we provided direct, global evidence that preferred synonymous codons are generally decoded more accurately than unpreferred codons. We found that relative translational accuracies of synonymous codons vary substantially among species, supporting the variable accuracy hypothesis. We obtained strong evidence that the ratio of cognate tRNA abundance to near-cognate tRNA abundance is a major determinant of a codon’s relative translational accuracy. Hence, the variable accuracies observed are mechanistically explained by the variation of the tRNA pool across species. These findings, together with the previous report on the selection for translational efficiency (*14*), suggest a model in which the tRNA pool and codon usage coevolve to improve both translational efficiency and accuracy (**Fig. 4a**). Specifically, mutation and drift can alter both codon frequencies and tRNA concentrations. The cellular translational efficiency is maximized when (transcriptomic) codon frequencies equal relative cognate tRNA concentrations (*14*), whereas the translational accuracy of a codon is maximized when the ratio of its cognate tRNA concentration to near-cognate tRNA concentration is maximized. Under this model, selections for translational efficiency and translational accuracy are related but not perfectly aligned, which could introduce tradeoffs between translational efficiency and accuracy (*27*). Indeed, our simulation of a simple genetic system with two amino acids, each encoded by two synonymous codons (**Fig. 4b**), found that imposing a selection for translational accuracy can lower translational efficiency (**Fig. 4c**).

**Figure 4.**
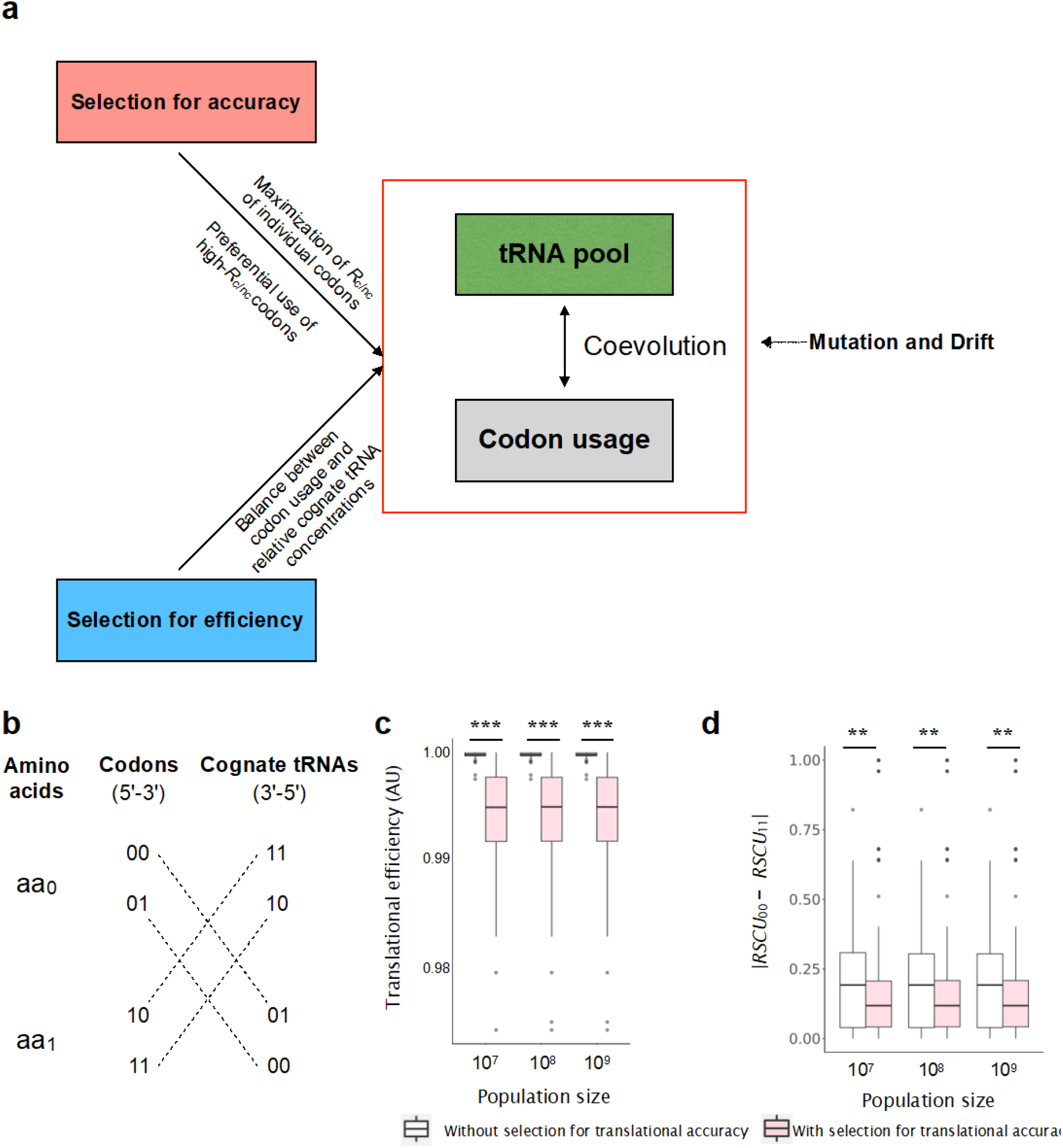
Selections for translational efficiency and accuracy shape the tRNA pool and codon usage. **a**, A model for the coevolution of the tRNA pool and codon usage driven by selections for translational efficiency and accuracy. **b**. A toy model with two amino acids, each encoded by two synonymous codons. A dotted line connects a codon with its near-cognate tRNA. **c**, Translational efficiency is significantly lower in the presence of selection for translational accuracy than in the absence of this selection. **d**, The absolute difference between the *RSCU* of 00 (*RSCU*_00_) and that of 11 (*RSCU*_11_) is smaller under the selection for translational accuracy than without this selection. With the selection, codon usage for aa_0_ and that for aa_1_ become coupled, because selection disfavors the cognate tRNA of the common codon of aa_0_ to become the near-cognate tRNA of the common codon of aa_1_, and vice versa. In c and d, each box plot shows the distribution from 200 replicates. The lower and higher edges of a box represent the first (qu_1_) and third (qu_3_) quartiles, respectively; the horizontal line inside the box indicates the median (md); the whiskers extend to the most extreme values inside inner fences, md ± 1.5(qu_3_-qu_1_); and the dots are outliers. ^*^, 0.01 ≤ *P* < 0.05, Wilcoxon rank-sum tests; ^**^, 0.001 ≤ *P* < 0.01; ^***^, *P* < 0.001.

Interestingly, our results imply that, even in the absence of selection for translational accuracy, the positive correlation between synonymous codon frequency and cognate tRNA concentration resulting from selection for translational efficiency (*14*) will likely render the cognate tRNA concentration relative to near-cognate tRNA concentration higher for more frequently used synonymous codons. Consequently, the positive correlation between the relative codon frequency and relative translational accuracy may arise in the absence of selection for translational accuracy. In fact, in *E. coli*, for 16 of the 18 amino acids with multiple synonymous codons, the codon with the highest cognate tRNA concentration has the highest *RR*_c/nc_. Upon shuffling the expression levels among tRNA species, we found that, for over one half of the 18 amino acids, the codon with the highest cognate tRNA concentration has the highest *RR*_c/nc_. This was true in each of 1000 shufflings. Nevertheless, in only 6 of these 1000 shufflings did all 18 amino acids exhibit the above feature. Thus, a high but non-perfect concordance between translational efficiency and accuracy is expected. Therefore, strictly speaking, the correlation in Fig. 1b by itself does not prove selection for translational accuracy. However, this correlation, in conjunction with the correlation between *RSCU* and *OR*, demonstrates that evolutionarily conserved sites tend to use preferred synonymous codons, which tend to be relatively accurately translated, hence proving the role of selection for translational accuracy in causing CUB, or the TAH. How codon usage and the tRNA pool evolve under the joint forces of selections for translational efficiency and accuracy in addition to mutation and drift is quite complex. For instance, because any tRNA is simultaneously a cognate tRNA for one or more codons and a near-cognate tRNA for some other codons, increasing the translational accuracy of a particular codon might be at the expense of the translational accuracy of another codon. Indeed, a previous study showed that artificially increasing the cognate tRNA expression levels for arginine codons can result in proteotoxic stress (*28*). This subtle tradeoff could cause non-independent uses of codons of different amino acids. This was indeed observed in the simulation aforementioned (**Fig. 4d)**. Future modeling work with realistic parameters might shed more light on this issue. In addition to impacting translational efficiency and accuracy, synonymous mutations also affect mRNA folding (*29*), mRNA stability (*30*), mRNA concentration (*30-32*), pre-mRNA splicing (*33*), and co-translational protein folding (*34, 35*), so additional selections may shape CUB and its evolution.

Our study has several caveats. First, in our calculation of a codon’s mistranslation rate, we lumped all mistranslations of the codon regardless of the erroneous amino acid it is translated to. Because different mistranslations of the same codon likely have differential fitness costs and because selection for translational accuracy likely minimizes the total fitness reduction caused by mistranslation instead of the mistranslation rate *per se*, properly weighting different mistranslations in *RMR* calculation will likely strengthen its correlation with *RSCU*. Second, when calculating the ratio of cognate tRNA concentration to near-cognate tRNA concentration, we did not consider the difference in interaction strength between different codons and anticodons (*36*). Future research that takes into account this interaction under physiological conditions may significantly improve the signal in the correlation analysis of Fig. 3. Third, our analysis in Fig. 3d was limited to taxa with >80 tRNA genes. Future research using tRNA expression levels (*26*) when they become available can confirm if the same pattern holds for taxa with fewer tRNA genes. Finally, due to data limitation, we did not consider tRNA expression variations across environments, cell cycle stages, or tissues (*37*). In the future, it would be interesting to study how such variations simultaneously impact translational efficiency and accuracy.

Our results might help design organisms with expanded code tables (*38*). Expanding the code table is realized by introducing unnatural tRNAs that are charged with non-canonical amino acids. The introduction of these tRNAs often leads to fitness defects due to mistranslation of normal codons (*39*). Our research suggests that one way to alleviate the proteotoxic stress is to identify potential near-cognate codons that could be mistranslated by the unnatural tRNA and adjust the natural tRNA pool to minimize the impact.

## MATERIALS AND METHODS

### Estimating relative mistranslation rates of synonymous codons from *E. coli* proteomic data

The proteomic data analyzed came from Table S1 in Modret *et al*. (*20*). The authors separately measured mistranslation events from high-solubility and low-solubility proteins using mass spectrometry, and both groups of events were considered in our analysis. We focused on the data from the wild-type strain BW25113 in the MOPS complete medium because (i) this dataset is the largest among datasets from all strain-medium combinations and (ii) no artificial perturbation such as mutation, drug, or amino acid depletion was applied (*20*). We first removed sites that cannot be traced to a unique original codon. We also filtered out sites showing an intensity of “NaN” for the unmodified (aka base) peptide or mistranslated (aka dependent) peptide. Because different synonymous codons tend to generate different mistranslations by mispairing with different near-cognate tRNAs, if these different mistranslations have different detection probabilities, the comparison between synonymous codons would be unfair. Unfortunately, some mistranslations produce mass shifts indistinguishable from post-translational modifications so cannot be reliably identified through mass spectrometry (*26*), which would produce exactly this situation in some cases. Therefore, we removed amino acids with undetectable mistranslations except for Leu and Ile. We kept these two amino acids because the only undetectable mistranslations for them are Leu to Ile and Ile to Leu, both can be considered benign due to the high physicochemical similarity between Leu and Ile (*40*). Considering the structure of the genetic code table, we found that the underestimation of the mistranslation rate due to the negligence of mistranslations between Leu and Ile is severer for unpreferred than preferred codons, suggesting that the actual strength of evidence for higher mistranslation rates of unpreferred than preferred synonymous codons is stronger than what is shown in Fig. 1. We then computed each codon’s absolute mistranslation rate by dividing the total intensity of mistranslated (i.e., dependent) peptides by that of all (i.e., dependent + base) peptides mapped to the codon. We divided each codon’s absolute mistranslation rate by the mean absolute mistranslation rate of all codons coding for the same amino acid to obtain the codon’s relative mistranslation rate (*RMR*). We removed amino acids without data for all of its synonymous codons because calculating *RMR* requires having data for all synonymous codons of an amino acid. In total, we computed *RMR* for 27 codons of 9 amino acids.

### Relative synonymous codon usage (*RSCU*), odds ratio (*OR*), and relative ratio of cognate tRNA concentration to near-cognate tRNA concentration (*RR*_c/nc_) for *E. coli*

Peptide and cDNA sequences of *E. coli* (genome assembly ASM584v2) and *S. enterica* (genome assembly ASM78381v1) were downloaded from Ensembl Bacteria (*41*). We computed *RSCU* of codon *j* of amino acid *i* from all coding sequences of *E. coli* by 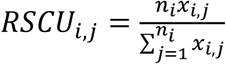 where *n*_i_ is the number of synonymous codons of amino acid *i* and *x*_*i,j*_ is the number of codon *j* of amino acid *i* in all coding sequences (*21*). Conventionally, *RSCU* is computed from highly expressed genes (*21*). However, due to the lack of gene expression information from most of the species analyzed, we computed *RSCU* from all genes. This should not qualitatively affect our analysis, because *RSCU* computed from highly expressed genes (e.g., the top 20% of genes) is nearly perfectly correlated with that computed from all genes (e.g., in *E. coli, r* = 0.96, *P* < 2.2×10^−16^).

To calculate the *OR* of each codon, we first identified one-to-one orthologous proteins between *E. coli* and *S. enterica* using OrthoFinder (*42*). Next, we aligned these one-to-one orthologs using MUSCLE (*43*), separating all amino acid sites into conserved and non-conserved sites. For a focal codon in gene *i*, we tabulated *a*_*i*_, number of times the focal codon is observed at conserved amino acid sites; *b*_*i*_, number of times the focal codon is observed at unconserved sites; *c*_*i*_, total number of times the focal codon’s synonymous codons are observed at conserved sites; and *d*_*i*_, total number of times the focal codon’s synonymous codons are observed at unconserved sites. Here, the focal codon’s synonymous codons do not include itself. *OR* for gene *i* equals (*a*_i_*d*_i_)/(*b*_i_*c*_i_). Using the Mantel-Haenszel procedure, we combined the odds ratios of the focal codon from individual genes into one odds ratio (*7*) by 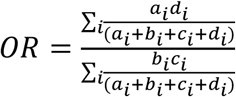.

To compute *RR*_c/nc_ of a codon, we tabulated the cognate tRNAs and near-cognate tRNAs of the codon. Cognate tRNAs are all tRNAs that can pair with the focal codon allowing wobble pairing at the 3rd codon position, while near-cognate tRNAs are tRNAs coded for a different amino acid but can pair with the focal codon with one base-pair mismatch (allowing wobble pairing at the 3rd codon position). We then weighted each tRNA by their average relative expression levels across three growth stages in the MOPS complete media (GEO number: GSE128812). Finally, we normalized the ratio for each codon by the average ratio of all codons coding for the same amino acid.

### *RSCU, OR*, and *RR*_c/nc_ for other species

*RSCU, OR*, and *RR*_c/nc_ were calculated for non-*E. coli* taxa as for *E. coli*, with the differences noted below. For the non-*E. coli* prokaryotic taxa, we downloaded the phylogenetic tree of 10,575 taxa from the Web of Life (*23*) (https://biocore.github.io/wol/) and identified sister taxa from the tree. Briefly, each pair of sister taxa are two taxa that are the single closest relative to each other in the tree. For each pair of sister taxa, we downloaded from the same web site their protein-coding DNA sequences, protein sequences, and tRNA gene copy number data. For eukaryotic model organisms, we downloaded protein-coding DNA sequences and protein sequences of human (*Homo sapiens*), mouse (*Mus musculus*), fly (*Drosophila melanogaster*), worm (*Caenorhabditis elegans*), and budding yeast (*Saccharomyces cerevisiae*) from the NCBI refseq database (*44*). We further downloaded the protein sequences of *Macaca mulatta* (as a relative of *H. sapiens*), *Rattus norvegicus* (as a relative of *M. musculus*), *Drosophila erecta* (as a relative of *D. melanogaster*), *Caenorhabditis briggsae* (as a relative of *C. elegans*), and *Saccharomyces paradoxus* (as a relative of *S. cerevisiae*) from the NCBI refseq database. The tRNA gene annotations in the five model organisms were downloaded from GtRNAdb (*24*). *RR*_c/nc_ was computed using tRNA gene copy numbers instead of tRNA expression levels.

### Statistical analysis

Many of the quantities estimated in our work, such as *RMR, RR*_c/nc_, *RSCU*, and *OR*, are not independent among synonymous codons. To deal with this non-independence in statistical tests, we applied permutation tests. Specifically, in **Fig. 1b**, we generated 1000 permuted samples by shuffling the absolute mistranslation rates among all codons and then re-estimated *RMR* values. We then computed the correlation between *RMR* and *RSCU* in each permuted sample while holding the *RSCU* value of each codon unchanged. *P* equals the fraction of permuted samples with the correlation coefficient more negative than that observed in the original sample. Similarly, when testing the correlation between *RMR* and *OR* (**Fig. 2b**), we shuffled the absolute mistranslation rate among all codons and recomputed *RMR* while holding the *OR* for each codon unchanged. When testing the correlation between *RMR* (or *OR*) and *RR*_c/nc_ (**Fig. 3**), we shuffled the absolute mistranslation rates among codons and the expression levels (or gene copy numbers) among tRNAs. Finally, when testing the correlation between *RMR* (or *RSCU*) and relative cognate tRNA concentration (**Fig. S4**), we shuffled the absolute mistranslation rate among codons and the expression level among tRNAs.

To estimate the standard error (SE) of the *RMR* of each codon, we constructed 1000 bootstrap samples by resampling the sites in the original data with replacement. Similarly, we estimated the SE of the *OR* of each codon by constructing 1000 bootstrapped *E. coli* genomes via resampling its genes that have one-to-one orthologs in *S. enterica*.

### Simulation

To assess the impact of selections for translational accuracy and efficiency on codon usage, we built a toy model with two amino acids: aa_0_ and aa_1_. Amino acid aa_0_ is encoded by synonymous codons 00 and 01 while aa_1_ is encoded by synonymous codons 10 and 11 (**Fig. 4b**). Codon-anticodon pairing follows the rule that 0 pairs with 1 and vice versa. The cognate tRNA of a codon has an anticodon that pairs perfectly with the codon, while the near-cognate tRNA has an anticodon that pairs with the codon with exactly one mismatch and carries the other amino acid.

We considered a unicellular organism with one gene consisting of *n* codons. We assumed that the mRNA level of the gene does not change in the evolution simulated and that ribosomes are in shortage. We defined the organismal fitness as follows.

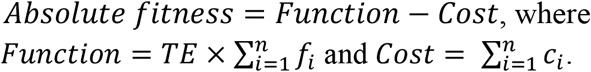

Here, *f*_i_ and *c*_i_ are the function and cost of codon *i*, respectively. We set *f*_i_ = *F*_i_ if codon *i* encodes the pre-specified optimal amino acid at the codon; otherwise, *f*_i_ = 0. For each *i, F*_i_ is a random variable sampled from an exponential distribution with the mean equal to 1 (*45*). Following a previous study (*14*), we set the expected codon selection time per amino acid aa_0_ as *t*_0_ = *p*_1_^2/^/*q*_1_+*p*_2_^2^/*q*_2_, where *p*_1_ and *p*_2_ = 1-*p*_1_ are the fractions of amino acid aa_0_ encoded by codon 00 and 01, respectively, and *q*_1_ and *q*_2_ =1-*q*_1_ are the fractions of corresponding cognate tRNAs among all tRNAs of aa_0_, respectively. We similarly set the expected codon selection time per amino acid aa_1_ and computed the total codon selection time of all codons. Translational efficiency *TE*, which is the number of codons translated per unit time, is the inverse of the total codon selection time. We set *c*_i_ = *TE* × *C*_i_ if codon *i* does not encode the pre-specified optimal amino acid at the codon; otherwise, 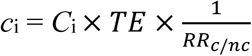. When there is no selection for translational accuracy, *C*_i_ = 0; otherwise, *C*_i_ for codon *i* is a random variable sampled from an exponential distribution with mean equal to 1. Note that *C*_i_ and *F*_i_ are independent from each other. *RR*_c/nc_ is computed as described in Results; the inverse of *RR*_c/nc_ measures the mistranslation rate.

We started the simulation with a coding sequence of 200 nucleotides, coding for 100 amino acids. Each site had a 50% chance to be 0 or 1. For simplicity, we assumed that the initial amino acid sequence is optimal such that the evolution in our simulation is largely about codon usage. For each of the four different tRNAs (with anticodons of 00, 01, 10, and 11, respectively), we sampled the initial copy number from 1, 2, and 3 with equal probabilities.

Next, we simulated the coevolution between the tRNA pool and codon usage following a strong selection, weak mutation regime. We first generate a mutation. With a probability of 0.02, it alters the copy number of a tRNA. In this case, we randomly pick a tRNA species to change its copy number by +1 or -1 with equal probabilities unless the copy number is 1, in which case it is +1. With a probability of 0.98, the mutation is a random point mutation at a randomly picked site of the coding sequence. The fitness of the mutant is then computed following the above fitness definition. The mutation is fixed with a probability of 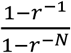, where *r* is the ratio of the absolute fitness of the mutant to that of the wild-type and *N* is the population size (*46*). The above mutation-selection process was repeated 100,000 rounds in each simulation to reach an equilibrium. For each *N*, we simulated 200 times with and 200 times without selection for translational accuracy.

## ACKNOWLEDGEMENTS

We thank Dr. Yitzhak Pilpel for consultation on the *E. coli* proteomic data in Mordret *et al*. (2019) and Wenfeng Qian, Jian-Rong Yang, and members of the Zhang laboratory for valuable comments. This work was supported by the U.S. National Institutes of Health research grant R35GM139484 to J.Z.

## Supplementary materials of

**Table S1.** Pearson (*r*) and Spearman (*ρ*) correlations Between *OR* and *RSCU* in eukaryotic model organisms.

| **Species** | **$r$** | **$P$** | **$\rho$** | **$P$** |
| --- | --- | --- | --- | --- |
| Fly | 0.84 | <0.001 | 0.83 | <0.001 |
| Human | 0.80 | <0.001 | 0.74 | <0.001 |
| Mouse | 0.74 | <0.001 | 0.68 | <0.001 |
| Worm | 0.42 | <0.001 | 0.39 | 0.002 |
| Yeast | 0.35 | 0.01 | 0.39 | 0.003 |

**Table S2.** Pearson (*r*) and Spearman (*ρ*) correlations between *OR* and *RR*_c/nc_ in eukaryotic model organisms.

| **Species** | **$r$** | **$P$** | **$\rho$** | **$P$** |
| --- | --- | --- | --- | --- |
| Fly | 0.21 | 0.109 | 0.25 | 0.046 |
| Human | 0.39 | 0.007 | 0.29 | 0.055 |
| Mouse | 0.29 | 0.081 | 0.41 | 0.018 |
| Worm | 0.54 | <0.001 | 0.57 | <0.001 |
| Yeast | 0.42 | 0.045 | 0.37 | 0.053 |

**Fig. S1.**
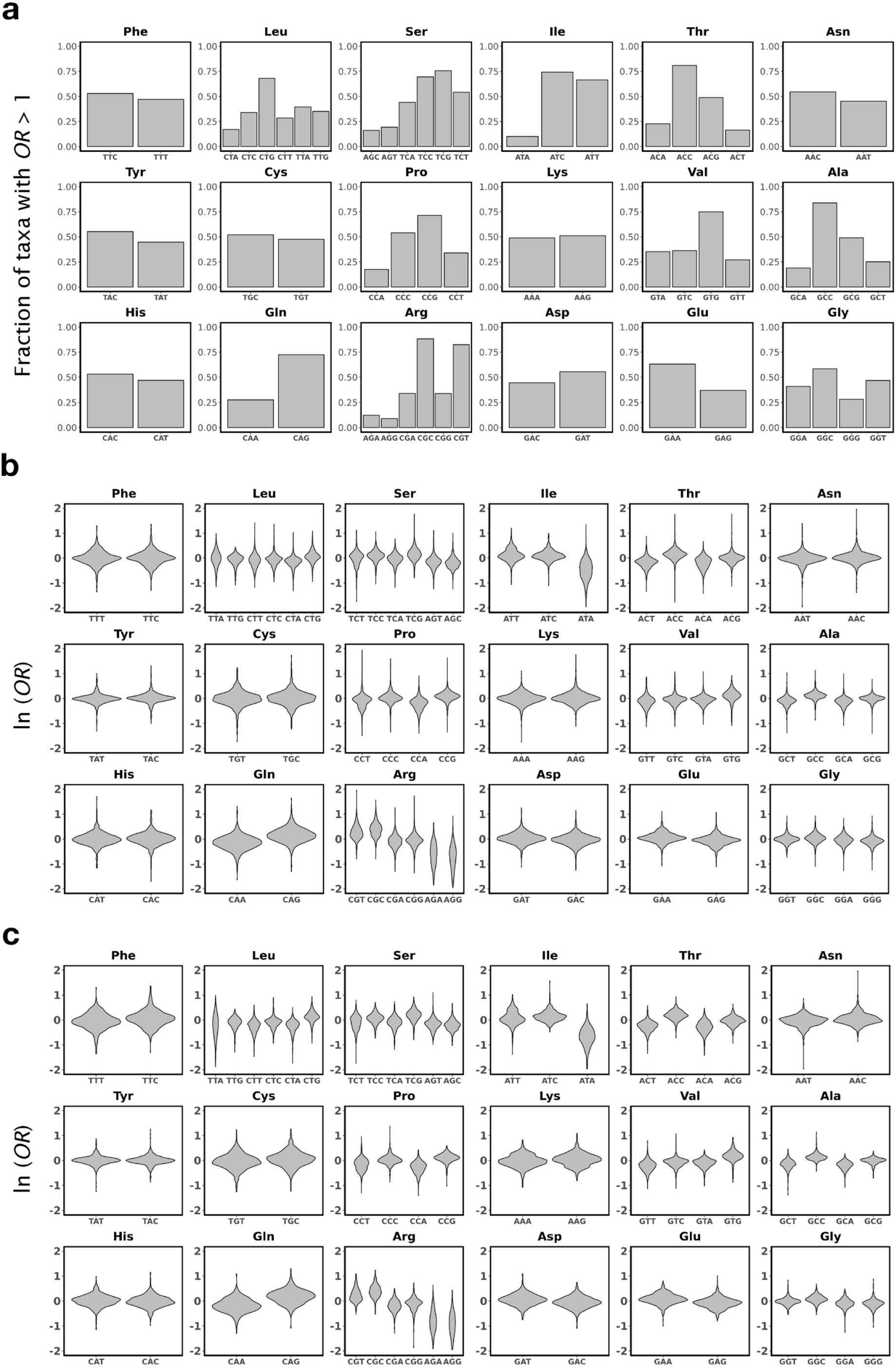
Observed patterns of variation of relative translational accuracies of synonymous codons across bacterial taxa are robust. **a**, Fraction of 1197 taxa with *OR* > 1 for each codon. **b**, Violin plots showing frequency distributions of ln(*OR*) of individual codons across a subset of taxa in which the number of occurrences of each synonymous codon considered in *OR* estimation is at least 1000. **c**, Violin plots showing frequency distributions of ln(*OR*) of individual codons across a subset of taxa with strong signals of selection for translational accuracy (i.e., Pearson’s correlation between *RSCU* and *OR* exceeds 0.5).

**Fig. S2.**
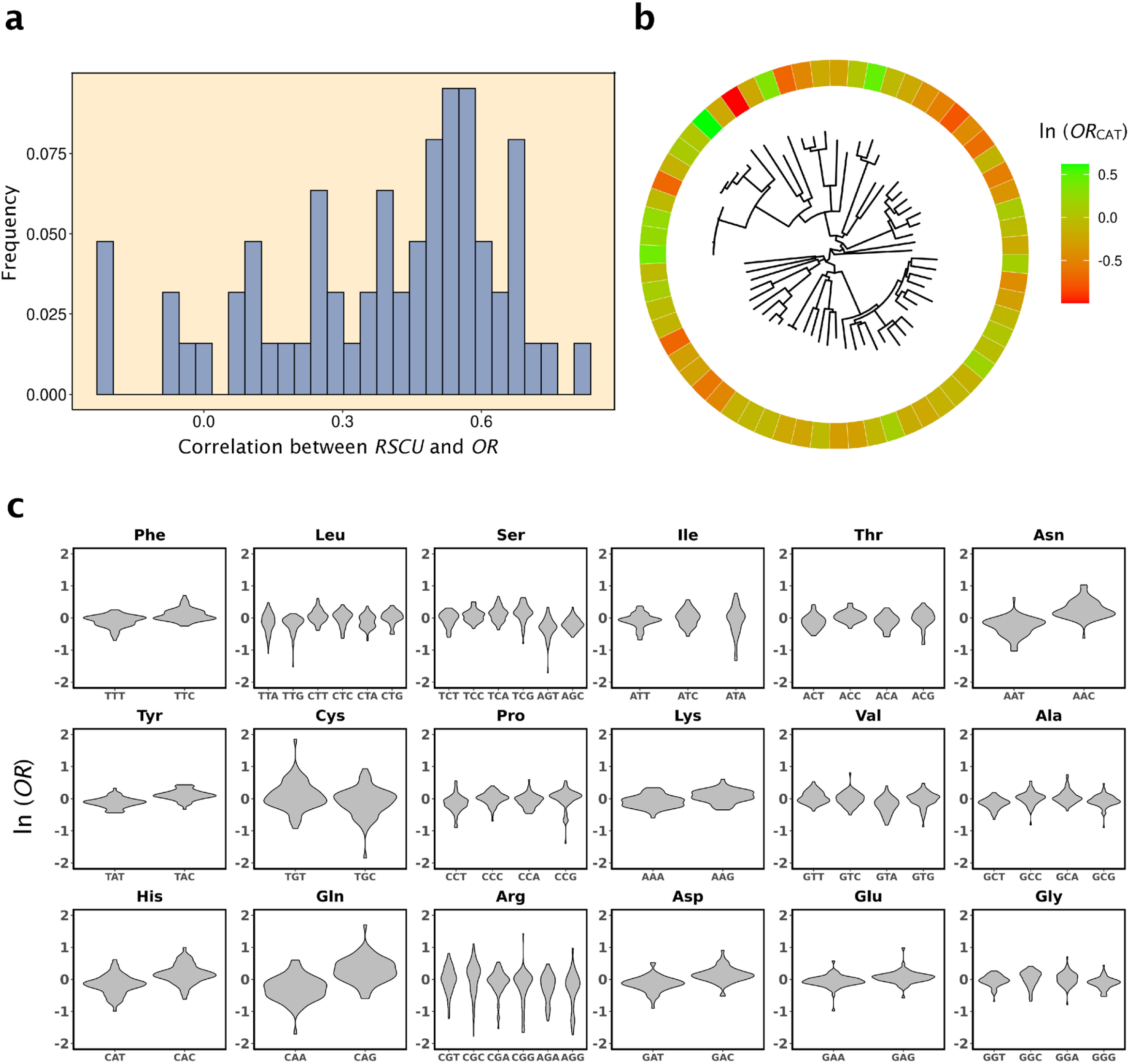
Variation of relative translational accuracies of synonymous codons among Archaea taxa. **a**, Frequency distribution of Pearson’s correlation between *RSCU* and *OR* in 63 taxa. Ninety percent of taxa show positive correlations. **b**, ln(*OR*) of codon CAT for each of the taxa arranged according to their phylogeny shown in the middle. **c**, Violin plots showing frequency distributions of ln(*OR*) of individual codons across taxa. ln(*OR*) appears less variable here than in Bacteria because of the much fewer Archaea taxa examined.

**Fig. S3.**
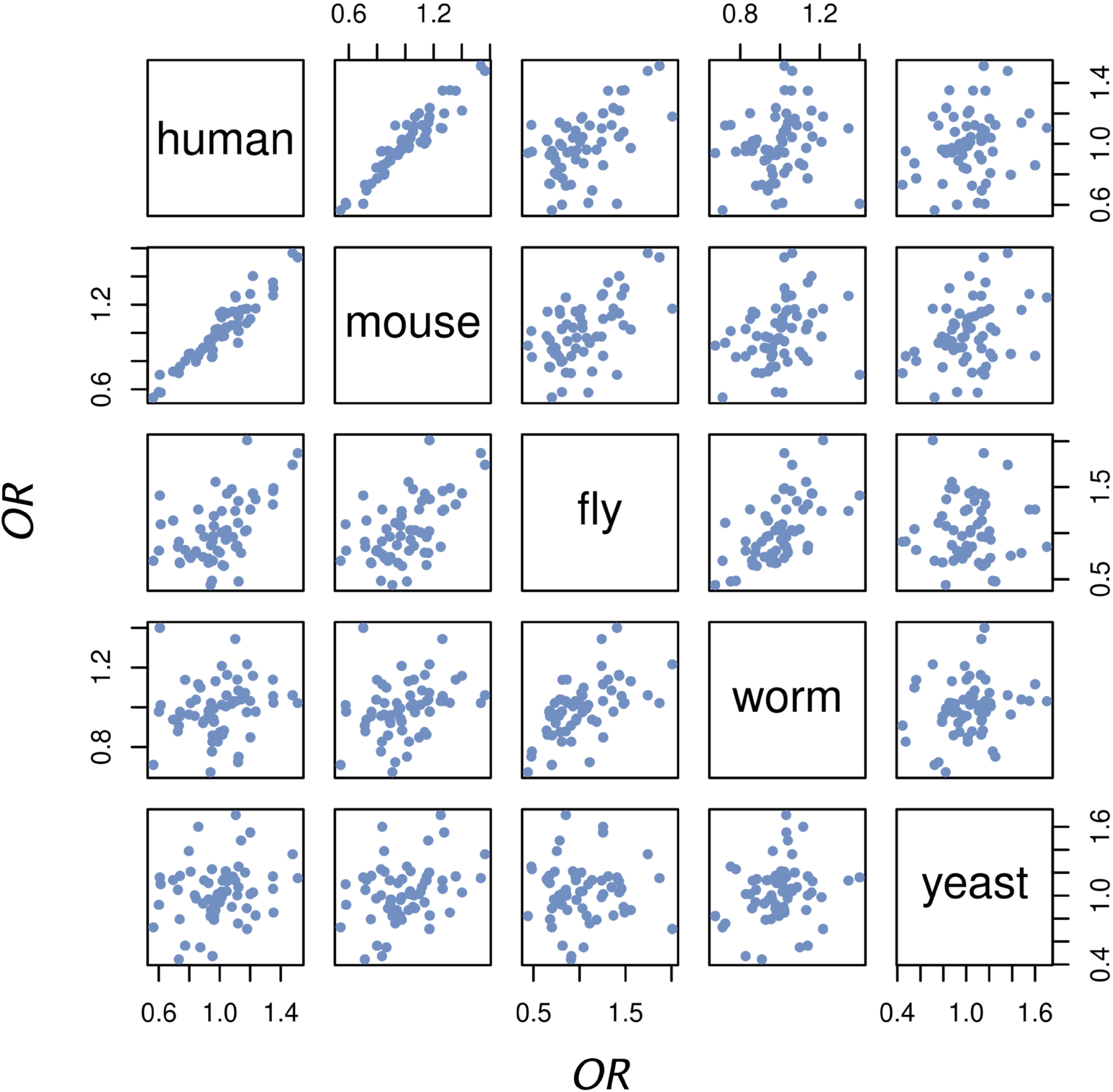
Correlation in odds ratio (*OR*) across codons between eukaryotic model organisms. Each dot is a codon.

**Fig. S4.**
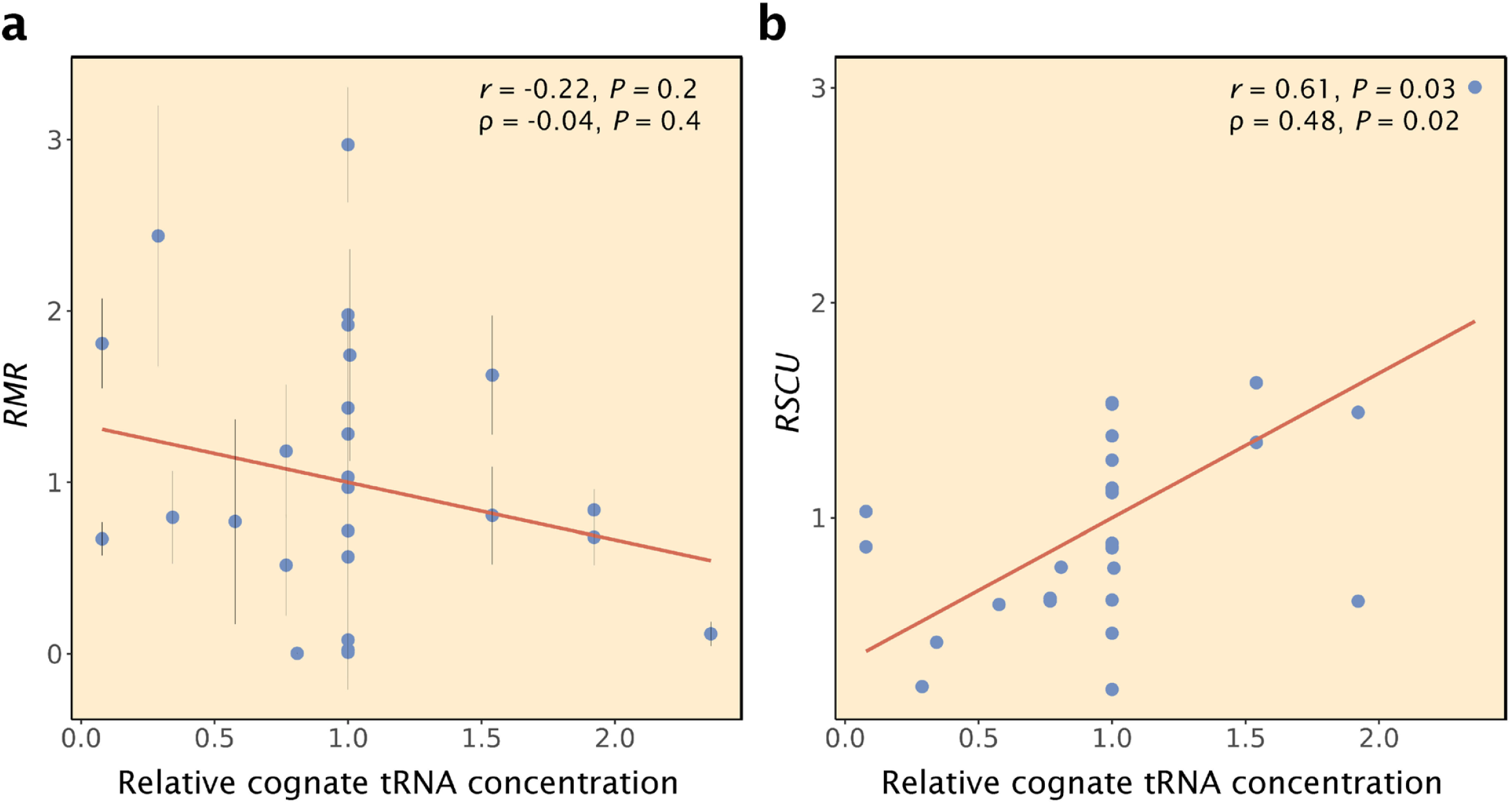
Relationship between the cognate tRNA concentration and *RMR* or *RSCU* in *E. coli*. **a**, *RMR* of a codon is not significantly correlated with its relative cognate tRNA concentration, which is its cognate tRNA concentration divided by the mean cognate tRNA concentration of all codons coding for the same amino acid. **b**, *RSCU* of a codon is significantly positively correlated with its relative cognate tRNA concentration. *P*-values are based on permutation tests. Only the 27 codons with *RMR* estimates are analyzed in each panel to allow a direct comparison.

